# Innate Immune sensing of Influenza A viral RNA through IFI16 promotes pyroptotic cell death

**DOI:** 10.1101/2021.02.13.431067

**Authors:** Shalabh Mishra, Athira S Raj, Akhilesh Kumar, Ashwathi Rajeevan, Puja Kumari, Himanshu Kumar

## Abstract

Programmed cell death pathways are triggered by various stresses or stimuli, including viral infections. The mechanism underlying the regulation of these pathways upon Influenza A virus IAV infection is not well characterized. We report that a cytosolic DNA sensor IFI16 is essential for the activation of programmed cell death pathways in IAV infected cells. We have identified that IFI16 functions as an RNA sensor for influenza A virus by binding to genomic RNA. The activation of IFI16 triggers the production of type I, III interferons, and also other pro-inflammatory cytokines via the STING-TBK1 and Pro-caspase-1 signaling axis, thereby promoting cell death (apoptosis and pyroptosis in IAV infected cells). Whereas, IFI16 knockdown cells showed reduced inflammatory responses and also prevented cell mortality during IAV infection. These results demonstrate the pivotal role of IFI16-mediated IAV sensing and its essential role in activating programmed cell death pathways.

## Introduction

The past century has witnessed several pandemics disrupting the socio-economic harmony of humankind. The influenza pandemic of 1917 and the current novel coronavirus pandemic are examples of the devastation caused by newly evolved viruses. The short replication time of viruses helps them acquire zoonotic potential through numerous mutations over a short period. Influenza is a substantial threat to public health as it caused multiple catastrophic pandemics killing millions of people around the world. It has a segmented genome and relatively high mutation rate, leading to better survivability(1) and evolvability(2). This eight segmented negative-sense single-stranded RNA virus from the Orthomyxoviridae family naturally infects diverse species, including birds and mammals. Mixing up or exchanging various genomic segments (antigenic shift) during co-infection in an intermediate host can result in the emergence of novel viral strains(3). Apart from the pandemics, seasonal Influenza virus infections account for more than 5 million cases annually, severely affecting children and older adults(4, 5). An effective therapeutic strategy against emerging influenza virus strains is perplexing because it mutates very fast and subverts host immunity and cellular machinery. However, novel therapeutic approaches targeting host factors, essential for establishing viral infection, can prove to be more effective.

The RNA genome of Influenza A virus (IAV) in infected cells is sensed by evolutionarily preserved germline-encoded pathogen recognition receptors (PRRs), including Toll-like Receptors (TLR) 3 and 7(6), Retinoic acid-inducible gene I (RIG-I)(7), NOD-like receptor family member NOD-, Leucine-rich repeat (LRR)-and pyrin domain-containing 3 (NLRP3)(8), and the Z-DNA binding protein 1 (ZBP1)(9) in distinct cellular compartments. Upon sensing, the PRRs elicit an array of signaling pathways leading to the innate antiviral state through robust production of type I, III interferons and pro-inflammatory cytokines via different transcription factors like nuclear factor kappa-light-chain-enhancer of activated B cells (NF-κB) and interferon regulatory factors (IRFs). Interferons further sensitize neighboring cells by inducing Interferon Stimulated Genes (ISGs) and collectively develop a potent antiviral state. In addition to this complex innate immune signaling cascade, interferons and pro-inflammatory cytokines also induce programmed cell death in virus-infected cells.

Programmed cell death is classified into various types based on the cues leading to cell death and macroscopic morphological variations. It has been observed in-vitro and in-vivo that the IAV induces apoptosis(10), primary necrosis(11), necroptosis(12), and pyroptosis(13, 14) in various cell types. Virus-associated programmed cell death was initially perceived as a host defense mechanism that limits viral replication by eliminating infected cells. However, recent studies indicate that IAV can manipulate host immunity to induce cell death, helping its propagation. The IAV proteins NS1(14–16), M1(17), PB1-F2(10, 18), and NP(19) have been reported to activate apoptotic pathways to evade inflammatory responses and defend their replicative niche. The types of cell death pathways elicited upon IAV infection are mostly known. However, the innate immune sensing and signaling pathways deciding the fate of the cell upon IAV infection remain poorly understood.

This study reports a novel role of well-known PRR Interferon Gamma Inducible protein (IFI) 16 in eliciting cell death in alveolar epithelial cells. IFI16 is an intracellular DNA sensor mediating TBK-1-dependent IFNβ production via an adaptor STING. One of the AIM2-like Receptor (ALR) family member, IFI16, contains an N-terminal Pyrin domain (PYD) and two C-terminal HIN domains that bind to DNA in a sequence-independent manner(20). IFI16 was thought to be a cytosolic sensor, but recent studies show that it contains a multipartite nuclear localization signal (NLS) and senses nucleic acid in cytoplasm and nucleus in a PAMP-localization-dependent manner(21). IFI16 plays a critical role during various DNA viruses (KSHV(22, 23), HSV-1(20), EBV(24), HCMV(25)), retrovirus (HIV(24)), and bacterial infections (*Listeria Monocytogenes*(26)) by restricting pathogens’ propagation. Several DNA virus proteins have evolved to inhibit or degrade IFI16, highlighting its vital role in defense against infections(25, 27). Recent studies show that IFI16 can also restrict various RNA virus infections (Sendai(28), EMCV(28), CHIKV(29)). However, the mechanism by which IFI16 defends against RNA viruses remains elusive, especially about IAV infection. Through high-throughput transcriptomic analysis after IFI16 knockdown, we uncovered the mechanistic insights about how IFI16 protects the host against the IAV. Our study shows that IFI16 restricts the IAV infection by sensing viral RNA (predominantly in the nucleus) and stimulating cell death.

## Materials and Methods

### Analysis of microarray data from the GEO database

Influenza H1N1 infected cell line and patient microarray data were obtained from the GEO database (GSE37571, GSE50628, GSE48466, GSE40844). Differentially expressed genes in the above datasets between mock-infected and H1N1 infected or Healthy and Patients were identified using the GEO2R online tool (10.1093/bioinformatics/btm254). Differentially expressed genes were plotted using various R packages.

### Cells, transfection, viruses, and reagents

A549 human alveolar basal epithelial cells (Cell Repository, NCCS, India) and HEK293T human embryonic kidney cells (ATCC CRL-3216) were cultured in Dulbecco’s modified Eagle’s medium (DMEM) supplemented with 10% Fetal Bovine Serum (FBS) and 1% penicillin-streptomycin. Small airway epithelial cells (SAECs; Lonza) were cultured and maintained according to the manufacturer’s instruction. Transfection of DNA and Poly(I-C) (InvivoGen) was performed with Lipofectamine 3000 (Invitrogen) in Opti-MEM as per the manufacturer’s protocol. Cells were infected in serum-free DMEM with the A/Puerto Rico/8/34 (PR8/H1N1), or NDV Lasota viruses at the MOIs mentioned in the figure legends. After one hour, cells were washed with Phosphate buffered saline (PBS) and replaced with DMEM containing 1% FBS. DMEM, FBS, Opti-MEM, and penicillin-streptomycin were purchased from Invitrogen (Carlsbad, CA, USA).

Anti-FLAG, anti-*β actin,* and anti-γ Tubulin antibodies were purchased from Sigma Aldrich (St. Louis, MO, USA). Anti-Caspase-3 and Anti caspase 1 antibodies were purchased from Cell Signaling Technology (Danvers, MA, USA). The anti-IFI16 antibody was purchased from Santa Cruz Biotechnology (Dallas, TX, USA). IR dye-labelled anti-Rabbit and anti-Mouse IgG (secondary antibody) were purchased from LI-COR.

### Short hairpin (sh)RNA mediated transient knock-down

shRNA clones were obtained from the whole RNAi human library for shRNA mediating silencing (Sigma, Aldrich) maintained at IISER, Bhopal, India. Cells were transfected with either control scrambled(scr) shRNA or specific shRNA clones against IFI16, IPS-1, STING, and MyD88 using Lipofectamine 3000(Invitrogen) in Opti-MEM as per the manufacturer’s protocol. The efficacy of each shRNA clone to downregulate the endogenous expression of IFI16, IPS-1, STING, and MyD88 in respective clones was measured by either semi-quantitative PCR or immunoblot.

### Generation of A/PR8/H1N1 virus

A/PR8/H1N1 viruses were generated using eight plasmid system as described previously(30–32).

### Cloning, Plasmids, and site-directed mutagenesis

pCMV3Tag1a (FLAG) was a kind gift from Professor Yan Yuan, University of Pennsylvania, Philadelphia. IFI16 plasmid was a kind gift from Professor Davide knipe. IFI16 was subcloned into PCMV 3tag1a for tagging with flag and in PEGFP N1 for tagging with GFP. The NLS mutant of IFI16 was made by site-directed mutagenesis using the primers 5′-GAGGCAGAAGGAAGTGGATGCTACTTCACC-3′ 5′-CCTTCTGCCTCTTTCTTGATAGGGCTGG-3′.

### Trypan Blue Exclusion Assay

Cells were treated stained with Trypan blue and counted stained (dead) and unstained (live)cells using a hemocytometer.

### Cell viability assay: MTT

[3-(4,5-dimethylthiazol-2-yl)-2,5-diphenyl tetrazolium bromide] assay was performed as described previously(33).

### Quantitative real-time reverse transcription-PCR

Total RNA was isolated using TRIzol reagent (Ambion/Invitrogen) and was used to prepare cDNA using iscript cDNA synthesis kit (Bio-Rad) following the manufacturer’s protocol. Gene expression was estimated by quantitative real-time PCR using SYBR green chemistry (Bio-Rad) and gene-specific primers18S (5′-CTGCTTTCCTCAACACCACA-3′ 5′-ATCCCTGAAAAGTTCCAGCA-3′) IFI16 (5′-ACTCCTGGAGCTCAGAACCC-3′ 5′-CTGTGTCTGTGTAGCCACTGT-3′) PR8 NP (5′-GGAGGGGTGAGAATGGACGA-3′ 5′-GTCCATACACACAGGCAGGC-3′) NDV (5′-GGAGGATGTTGGCAGCATT-3′ 5′-GTCAACATATACACCTCATC-3′) IFNβ (5′-AGCTGCAGCAGTTCCAGAAG-3′ 5′-AGTCTCATTCCAGCCAGTGC-3′ IL-6 (5′-CTCAGCCCTGAGAAAGGAGA-3′ 5′-CCAGGCAAGTCTCCTCATTG-3′) IP-10 (5′-TGGCATTCAAGGAGGTACCTCTC-3′ 5′-TGATCTCAACACGTGGACAAA-3′) CASP8 (5′-AGAGTCTGTGCCCAAATCAAC-3′ 5′-GCTGCTTCTCTCTTTGCTGAA-3′) BAX (5′-TCCCCCCGAGAGGTCTTTT-3′ 5′-CGGCCCCAGTTGAAGTTG-3′) BAK (5′-CATCAACCGACGCTATGACTC-3′ 5′-GTCAGGCCATGCTGGTAGAC-3′) PRKCE (5′-CAACGGACGCAAGATCGAG-3′ 5′-CTGGCTCCAGATCAATCCAGT-3′) XIAP (5′-TTTGCCTTAGACAGGCCATC-3′ 5′-TTTCCACCACAACAAAAGCA-3′) Survivin (5′-AGAACTGGCCCTTCTTGGAGG-3′ 5′-CTTTTTATGTTCCTCTATGGGGTC-3′) cIAP1(5′-AGCTAGTCTGGGATCCACCTC-3′ 5′-GGGGTTAGTCCTCGATGAAG-3′) TRAIL (5′-AGCAATGCCACTTTTGGAGT-3′ 5′-TTCACAGTGCTCCTGCAGTC-3′)

### Fluorescence-activated cell sorting Cytometry Analysis

Cells were stained with FITC labeled Annexin V and propidium iodide (Invitrogen) based on the manufacturer’s instructions. Stained cells were analysed using a FACS Aria III (Becton Dickinson), and data were analysed by using FlowJo software (FlowJo, Ashland, OR, USA).

### RNA-Seq analysis

Cells were harvested in TRIzol; total RNA was isolated and assessed for quality. cDNA libraries were prepared using TruSeq technology according to the manufacturer’s protocol (Illumina, San Diego, CA). Libraries were sequenced using NextSeq500 with a read length (2 × 75 bp) by Eurofins Genomic India Private Limited, India. FastQC was used to assess the read quality of raw data (http://www.bioinformatics.babraham.ac.uk/projects/fastqc/). Trimmomatic was used to remove the Illumina adaptors and filter the reads using a sliding window approach(34). Approximately 20 million cleaned pair-end sequencing reads from each sample were uploaded to the Galaxy web platform and were analysed at https://usegalaxy.org. HISAT2 was used to map the reads with the reference human genome (hg38). Aligned RNA seq reads were assembled to transcripts, and abundance was quantified by using StringTie. Differential expression analysis of genes between groups was done using DESeq2(35). Various R packages were used to visualize the expression and differential expression outcomes. Gene ontology (GO) analysis was done using the web-based Gene Set Analysis toolkit, and analysis of upregulated KEGG pathways was done using Enrichr. Cluster 3.0 and TreeView 1.1.6 were used for making heat maps. All the addressed analysis was demonstrated as described previously(36).

### Enzyme-linked immunosorbent assay (ELISA)

Scrambled (shSCR) or IFI16(shIFI16) knocked down A549 were infected with PR8 or NDV, and culture supernatants were collected 36 to 40hr post-infection and analysed for IL-6 and IP-10 cytokines according to the manufacture’s protocol (Becton Dickinson)

### Western blotting analysis

For endogenous detection of protein, scrambled (shSCR) or IFI16 (shIFI16) knocked down A549 cells (grown in a 6well plate) were lysed in 30 μl of standard cell lysis buffer, and ~ 8 μg of protein was loaded to each well. Immunoblotting was done as described previously using anti-*β* actin, anti-γ Tubulin, anti-Caspase3, anti-Caspase1, anti-flag, and anti-IFI16 antibody(37).

### ImageJ Image analysis

For western blot densitometry, in the immunoblots, an area of interest was selected on the smeary part above the bands, and the mean density of the selected area was calculated.

### RNA immunoprecipitation

Cells infected with PR8 24 hours post-transfection were harvested using standard lysis buffer supplemented with 1x protease inhibitor cocktail (Sigma Aldrich). 1/10 of the lysate was taken as input control. Immunoprecipitation was done using M2 flag affinity beads (Sigma) by incubating overnight with the lysate followed by multiple washes as per the manufacturer’s instruction. The RNA was isolated from the input, and the immunoprecipitated fraction using the TRIzol reagent and cDNA was made. Further, the pull-down viral RNA expression was quantified using qPCR, as described previously(9).

### Caspase-1 Assay

Caspase 1 activity was determined using FAM-FLICA caspase 1 assay kit (Immunochemistry Technologies) following the manufacturer’s protocol.

### Statistical analysis

All the experiments were done with appropriate controls, as mentioned in the legends. Experiments were carried out with triplicates or duplicates at least three times independently. GraphPad Prism 5.0 (GraphPad Software, La Jolla, CA, USA) was used for statistical analysis and plotting Graphs. Differences between the two groups were compared by an unpaired, two-tailed Student’s t-test, and differences between three groups or more were compared by ANOVA with the Newman-Keuls test. Differences were considered to be statistically significant when P-value P < 0.05. Statistical significance in the figures is indicated as follows: ***P < 0.001, **P < 0.01, *P < 0.05; ns, not significant

### Data availability

RNA-Seq data have been submitted to the Gene Expression Omnibus (NCBI-GEO) database under accession number (GSE163705)

## Results

### IAV induces cell death in IFI16 dependent manner

To decipher the role of RNA virus-sensing pathways in IAV induced cell death, we performed shRNA mediated transient knockdown of TLR adaptor MyD88 and the RLR adaptor IPS-1, and the adaptor for DNA-sensing pathway STING in human alveolar epithelial (A549) cells followed by Influenza A Virus (IAV) A/Puerto Rico/8/34 (PR8/H1N1) infection (**Supplementary figure1A**). Cells with STING knockdown showed the most resistance to IAV-induced cell death (**Supplementary figure1B**). Additionally, it has been previously reported that cells lacking the IFN-receptor were fully resistant to IAV-induced cell death(38). This suggests that IFN-mediated signaling plays an essential role in cell death upon IAV infection. In order to identify IFN-signaling genes that may have a possible role in IAV-mediated cell death, we re-analysed publicly available transcriptomic data from the NCBI-GEO database obtained from A549. Comparative analysis of uninfected and IAV-infected cells revealed a significant increase in the expression of nucleic-acid sensors and pro-inflammatory cytokines.

Interestingly, one of the most consistently upregulated nucleic-acid sensors in IAV-infected samples was the gene IFI16 (Interferon Gamma Inducible Protein 16) **(Supplementary figure 1C).** IFI16 has previously been identified as a cytosolic-double-stranded (ds) DNA sensor, which initiates type-I interferon dependent responses. Thus, IFI16 was of potential interest because its role in regulating the Influenza virus is not known, an RNA virus, and inducing cell death.

To probe the role of IFI16 in IAV induced cell death, IFI16 was knocked down in A549 cells using gene-specific shRNA **(Supplementary figure 2A),** and cell death in IAV-infected cells was analysed by MTT assay, Trypan-blue exclusion, propidium iodide (PI), and also by Annexin-PI FACS. We found that IFI16 knockdown cells were more than 80% resistant to IAV-induced cell death **(Figure 1A, 1B, 1C, 1D, Supplementary figure 2B).** Notably, IFI16 knockdown cells were resistant to cell death induced by IAV but not by another RNA virus, Newcastle Disease virus (NDV), suggesting an IAV-specific, IFI16-dependent cell death. Taken together, these data suggest a critical role of IFI16 in the regulation of cell death/survival during IAV infection.

**Figure 1.**
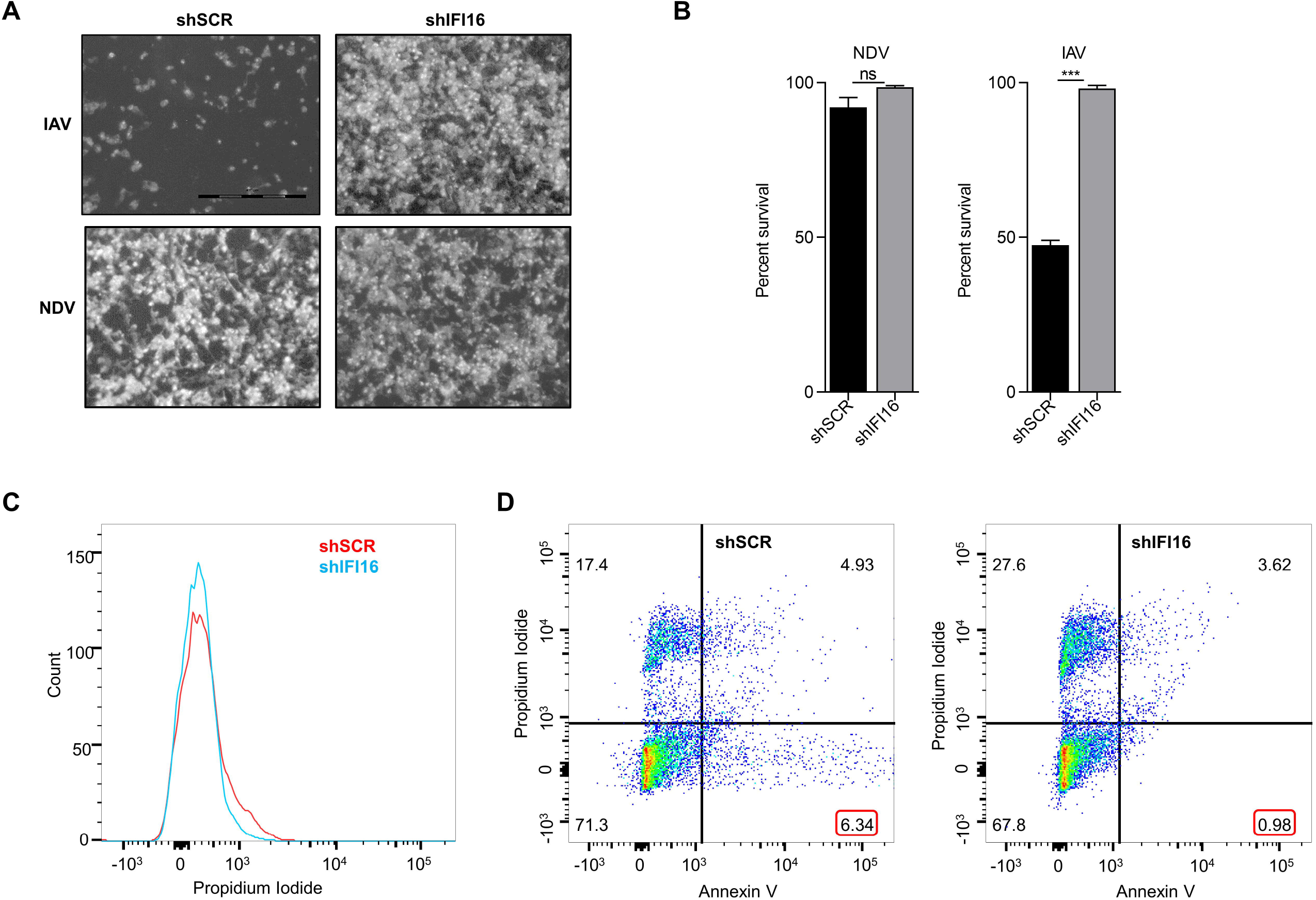
IAV induces cell death in IFI16 dependent manner. (A) A549 cells were transiently transfected with 2.0 μg either shRNA of IFI16 (shIFI16) or nonspecific scrambled control (shSCR) for 48 h then infected with IAV (MOI 10) (Upper panel) and NDV (MOI 2) (lower panel). 24 h post images were taken using 10x objective lens on an inverted microscope, and the cell viability was determined using the MTT assay. (B) and (C) 24 h post IAV infection, cells were stained with Propidium Iodide (PI) alone and PI + annexin V/, and apoptosis was determined by using flow cytometric analysis.

### IFI16 induce programmed cell death in IAV infected cells

To gain mechanistic insights into IFI16-mediate cell death, we performed whole transcriptome analysis of A549 cells transfected either with shSCR or shIFI16, followed by infection with IAV for 24 h, and subjected to RNA-Seq analysis in duplicates as shown in the schematic **(Figure 2A**). Principal component analysis of the gene expression demonstrates that samples cluster in distinct groups according to their treatment **(Figure 2B).** Differential gene expression analysis revealed that 947 genes were significantly upregulated, while 822 were downregulated **(Figure 2C).** Moreover, pathway analysis of significantly dysregulated genes using different tools indicated that IFI16 knockdown leads to the downregulation of interferon signaling pathways, TNF and NF-kB signaling, and caspase-mediated apoptosis signaling pathways **(Figure 2D).** We noticed that a large number of genes associated with apoptosis pathway were dysregulated **(Figure 2E).** The mRNA expression of selected pro-and anti-apoptotic genes was again confirmed by qRT-PCR **(Figure 2F).** These results suggest that IAV induced cell death is mediated through type I interferons, pro-inflammatory cytokines, and caspases and is IFI16-dependent.

**Figure 2.**
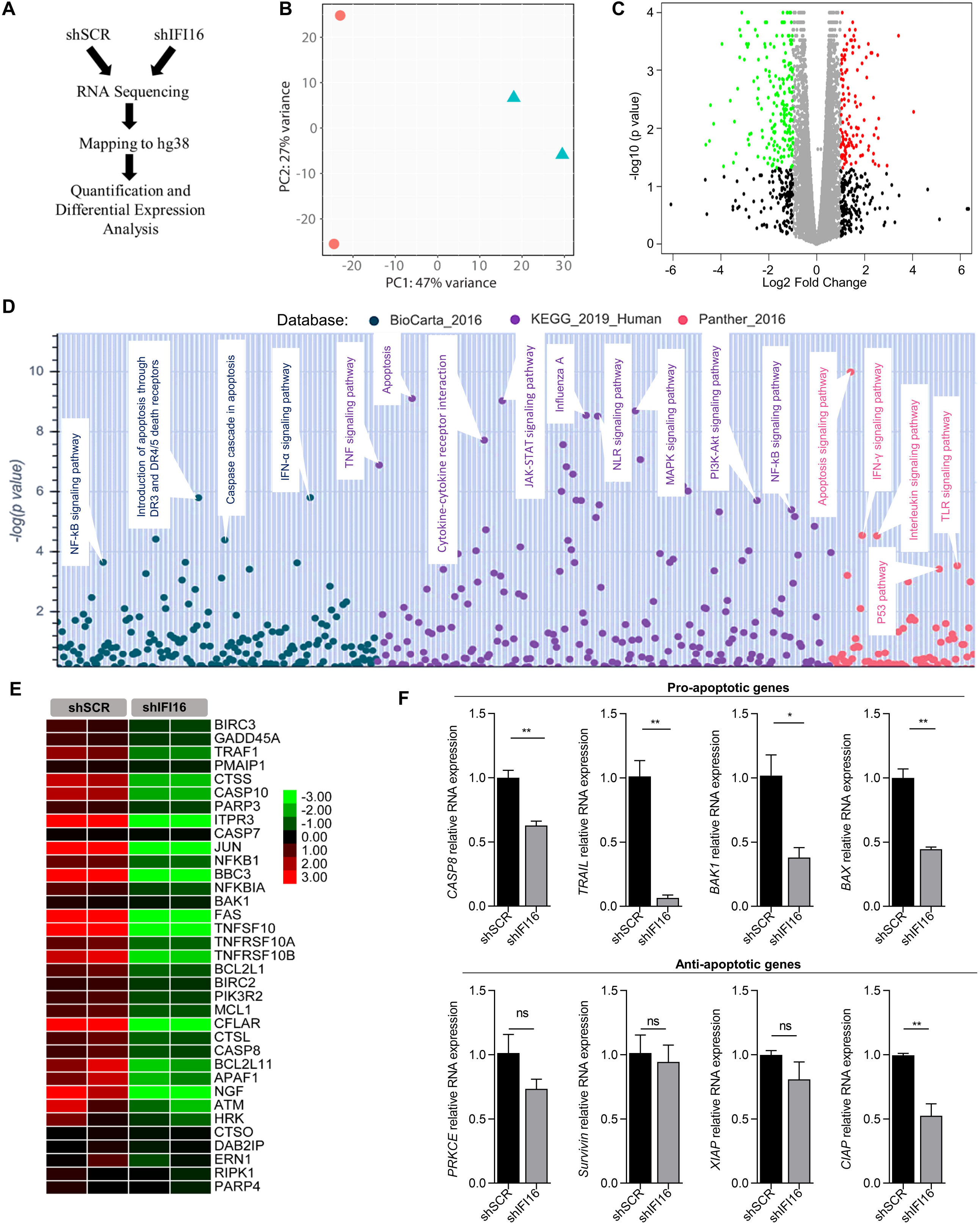
IFI16 induce programmed cell death in IAV infected cells. (A) A schematic representation of RNA-Sequencing data analysis. (B) Principal component analysis of samples resulted in the formation of two distinct groups (shSCR and shIFI16, in duplicates) according to their treatment. (C) Volcano plot showing differentially expressed genes in shIFI16 group. (D) Pathway analysis of significantly upregulated genes using Biocarta_16, KEGG_2019_human, and Panther_2016 database. (E) Heatmap showing expression of genes associated with apoptosis pathway in different samples. (F) mRNA expression of selected pro-and anti-apoptotic genes was again confirmed by qRT-PCR.

### IFI16 induces type I interferon and pro-inflammatory cytokines during IAV infection

We noticed that apart from upregulating apoptotic genes, knocking down of IFI16 also downregulates interferon signaling and pro-inflammatory cytokine production pathways during IAV infection **(Figure 3A, 3B, and Supplementary figure 3A)**. Previous studies have shown that IAV infection activates apoptosis and pyroptosis by triggering PRR induced gene expression of pro-inflammatory cytokines and type I interferons and type I interferons-inducible genes. Therefore, to further investigate which of these pathways are involved in IAV induced cell death, we used pharmacological inhibitors to pyroptosis and necroptosis pathways, Z-VAD-FMK, a pan-caspase inhibitor, and necrostatin-1, respectively. Only Z-VAD-FMK but not necrostatin-1 could inhibit cell death, indicating that IAV induces cell death through pyroptosis **(Figure 3C**). Also, only Z-VAD-FMK was able to rescue the cell death induced by IFI16 knockdown (**Figure 3D**), indicating IAV activates pyroptotic cell death in an IFI16 dependent manner. Furthermore, expression of the pyroptosis inducing cysteine proteases caspase 1 and caspase 3 was significantly reduced in shIFI16 cells compared to shSCR **(Figure 3E).** Activation of caspase-1 was analysed by FAM-FLICA caspase-1 assay and was found to be less in shIFI16 cells, which confirms the requirement of IFI16 and type-I interferon signaling in the activation of pyroptosis during IAV infection **(Figure 3F)**. We found that this phenomenon is specific to IAV, and IFI16 is dispensable for activating and releasing cytokines in response to other RNA virus infection (NDV) (**Supplementary figure 3B, 3C, 3D**). These data suggest a unique, type-I interferon and IFI16 dependent pathway of pyroptosis activation upon IAV infection, which was not previously reported.

**Figure 3.**
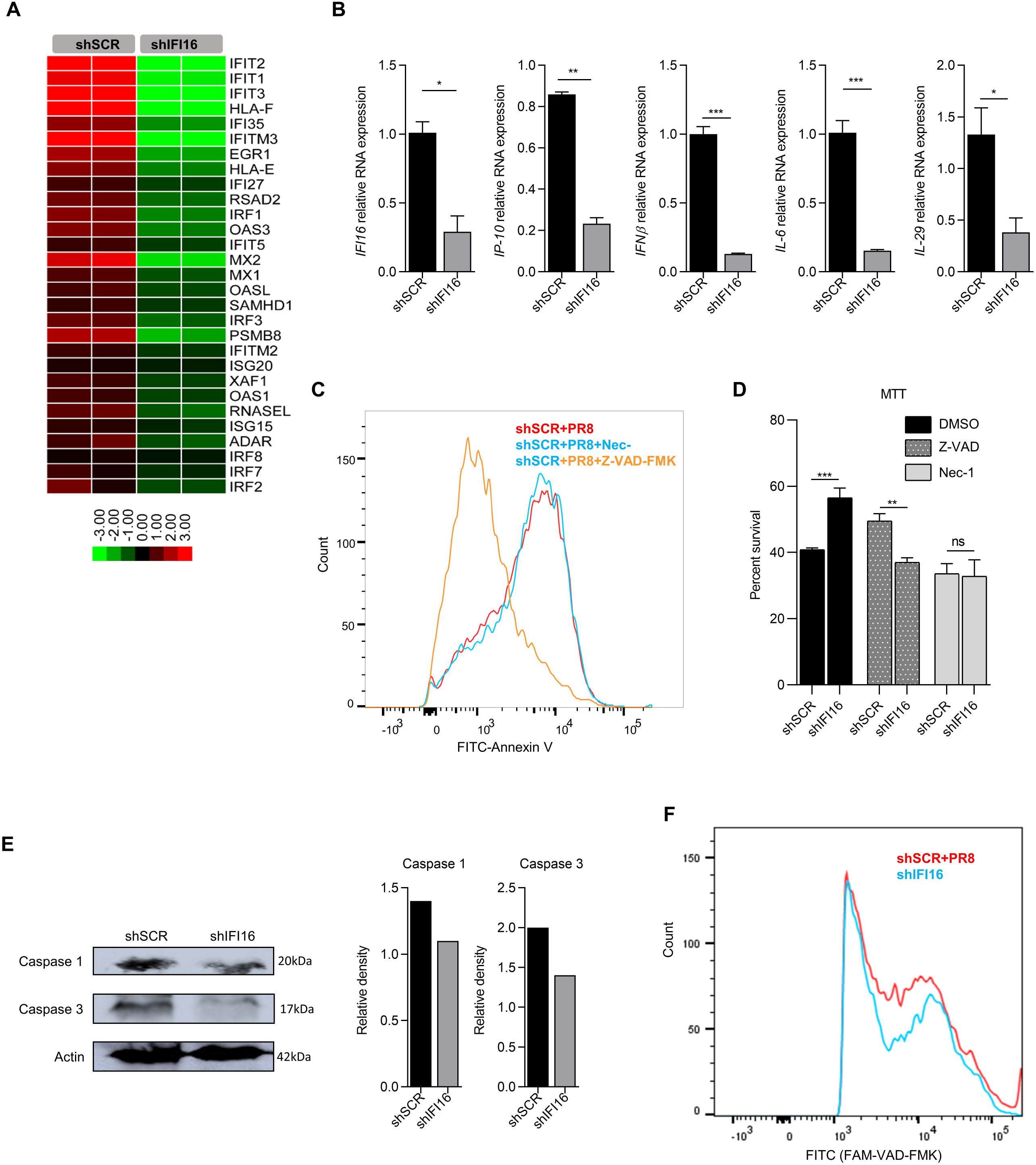
IFI16 induces type I interferon and pro-inflammatory cytokines during IAV infection. (A) Heatmap showing expression of interferon stimulated genes (ISGs) during IAV infection in different samples. (B) mRNA expression of interferons, ISGs and pro-inflammatory cytokines was again confirmed by qRT-PCR. (C) A549 cells were transiently transfected with 2.0 μg of scrambled control(shSCR) for 48 h. Cells were treated with DMSO, Z-VAD FMK (Z-VAD) or Necrostatin-1(Nec-1) one hour prior to infection and then infected with IAV (MOI 10) for 24 h. Cells were then stained with annexin V/PI, and analysed using flow cytometry for determination of apoptosis. (D) Cells were treated with Z-VAD FMK or Nec-1 in shSCR and shIFI16 groups prior to IAV infection. The cell viability was determined using the MTT assay. (E) Western blot analysis of Caspase 1 and Caspase 3 in shSCR and shIFI16 groups post IAV infection. (F) FAM-FLICA caspase 1 assay was performed to check the activation of caspase 1, and analysis was done through flow cytometry.

### Cell death induced by IAV depends on replication of viral genomic RNA

Programmed cell death is either initiated by extrinsic (cytokines) or intrinsic (TRAIL, p53) pathways. To determine the main driving factor for cell death, cytokines, or viral RNA replication, we examined the potential paracrine-signaling effects on cell death induction in uninfected cells using a transwell culture system. As shown in the representative figure **(Figure 4A),** cells were either mock-infected in both compartments or were infected with IAV only in the top compartment; the transwell culture system only allows paracrine signaling factors to move across. Upper compartments were removed 24 hours post-infection, and cell viability in the lower compartment was measured using MTT assay. Cells infected in the top compartment did not induce cell death in uninfected cells in the lower compartment, indicating that paracrine-signaling is not associated with IAV-induced cell death. Conversely, pre-treatment of infected cells with the RNA polymerase II and viral RNA-dependent RNA polymerase (RdRp) inhibitor actinomycin-D suppressed the cell death **(Figure 4B),** and UV irradiation of IAV before infection also significantly reduced cell death **(Figure 4C)** in proportion to the reduction in the virus infectivity. Altogether, these results suggest that cell death induction involves both extrinsic and intrinsic mechanisms and likely requires virus RNA replication.

**Figure 4.**
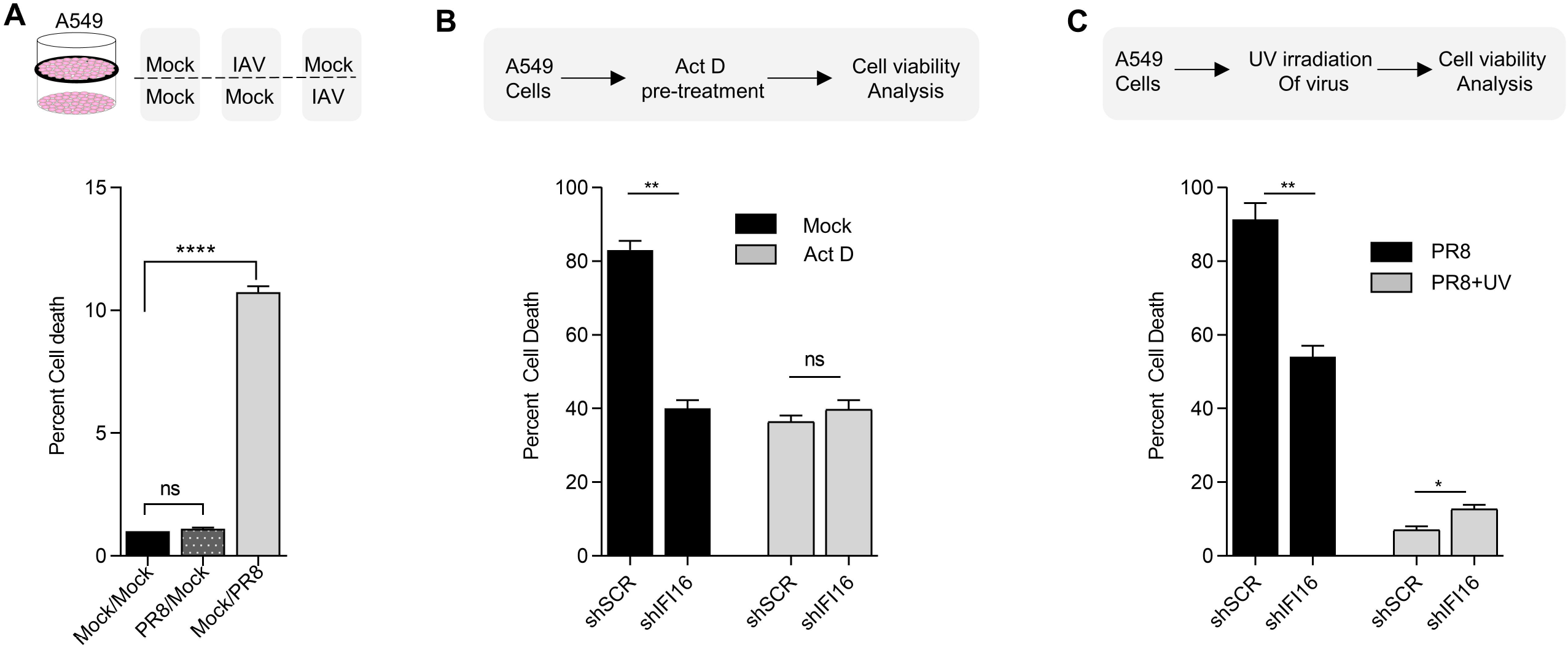
Cell death induced by IAV depends on replication of viral genomic RNA. (A) Transwell experiment, where mock-infected A549 (lower chamber) were exposed to paracrine signaling from mock-or IAV-infected A549 (upper chamber). IAV-infected cells in the lower chamber were used as a positive control. Cells were then stained with annexin V/PI, and apoptosis was determined by using flow cytometry. (B) 30-min pre-treatment was done in A549 cells (shSCR and shIFI16) with actinomycin D (Act D) (2 μM) before IAV infection. The percentage of cell death was calculated using the Trypan blue exclusion assay. (C) Infectivity of IAV was reduced after UV irradiation of virus. Both IAV and IAV+UV were used to infect A549 cells (shSCR and shIFI16). The percentage of cell death was calculated using the Trypan blue exclusion assay.

### IFI16 binds to Influenza virus RNA restricting its replication

IFI16 is known to sense cytosolic DNA from various DNA viruses and intracellular bacteria. However, it is not clear if it plays any role during IAV infection. To find if IFI16 interacts with IAV RNA, we performed RNA-immunoprecipitation (RNA-IP) of Flag-tagged IFI16 protein using anti-flag antibody, viral RNA in the precipitate was analysed by RT-PCR. IAV RNA was present in the anti-Flag pulldown, and interestingly, the enrichment of IAV RNA was orders of magnitude greater than of RIG-I, a well-known sensor of IAV, despite similar levels of IAV RNA in the input **(Figure 5A)**.

**Figure 5.**
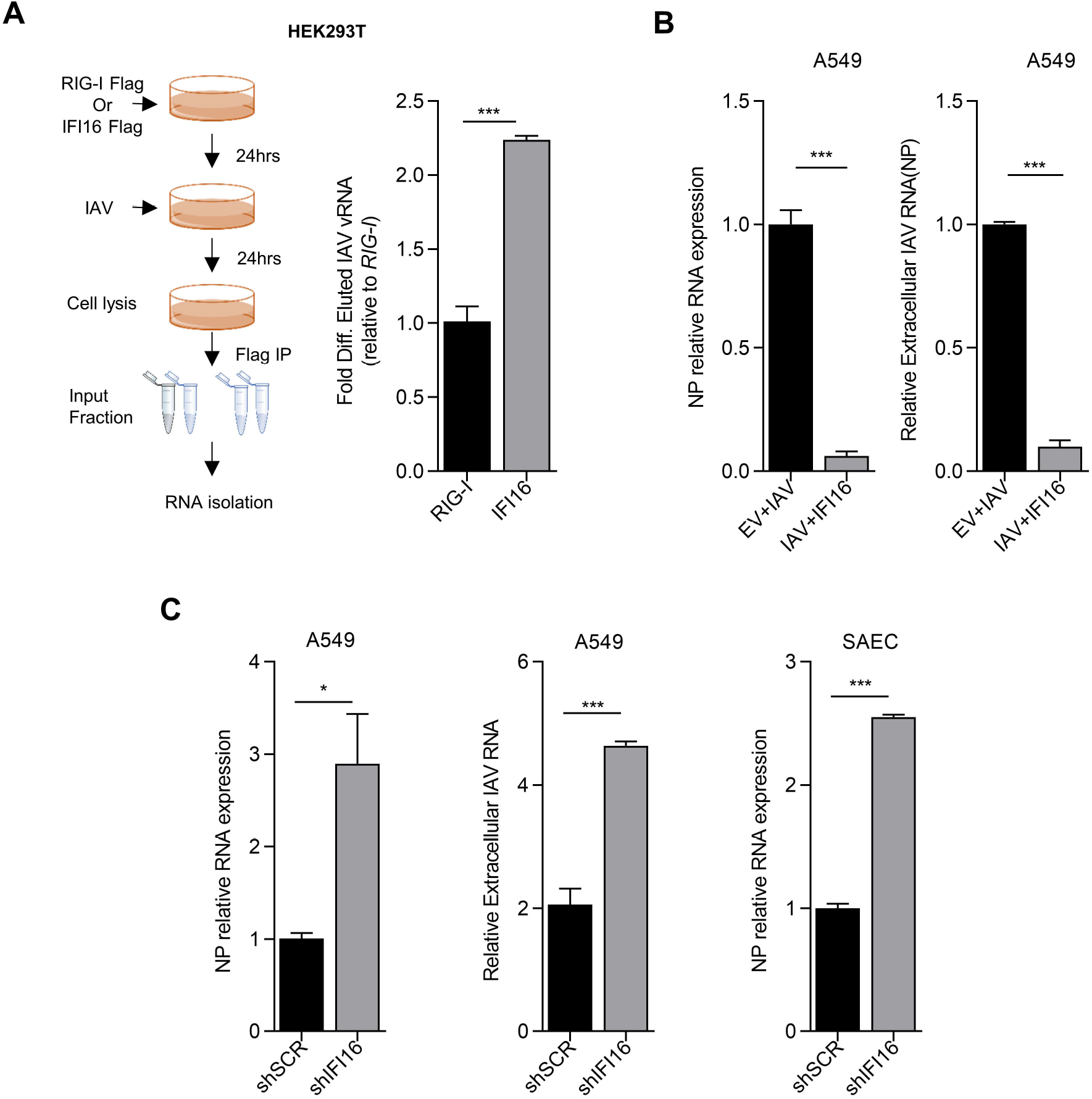
IFI16 binds to Influenza virus RNA restricting its replication. (A) Schematic for RNA immunoprecipitation (RNA-IP) (left side). 2.0 μg of Flag-tagged IFI16 and RIG-I were overexpressed in HEK293T cells and then infected with IAV (MOI 10) for 24h. RNA-IP was performed using Anti-flag M2 Affinity gel, and viral RNA in the precipitate was analysed by qRT-PCR using Input RNA for normalization. (B) A549 cells were transfected with plasmid expressing IFI16 or the empty vector backbone (EV) and infected with IAV (MOI 10) for 24 h. Relative NP RNA was measured in total RNA (Left panel) and extracellular RNA (Right panel) by qRT-PCR. NP RNA expression was normalized to mock infected sample. (C) A549 and SAEC cells were transiently transfected with 2.0 μg of shIFI16 or scrambled control plasmids for 48 h then infected with IAV (MOI 10). Relative NP RNA in the total RNA was analysed by qRT-PCR.

Next, we asked if binding of IFI16 to viral RNA also affects IAV replication; overexpression of IFI16 led to a decrease in IAV RNA inside cells as well as in supernatant **(Figure 5B).** Similarly, Knockdown of IFI16 led to an increase in IAV infection **(Figure 5C)** in both A549 and in primary small airway epithelial cells (SAEC). Besides, levels of pro-inflammatory cytokines also decreased in IFI16 knocked-down cells infected with IAV.

### Nuclear localization of IFI16 is essential in restraining infection and sensing IAV genomic RNA

To test the effect of IFI16’s subcellular localization on its viral RNA sensing, we mutated nuclear localization sequences (NLS) on the N-terminus of the IFI16 (**Figure 6A**). NLS mutation was able to restrict most of the (>70%) IFI16 protein to the cytosol (**Figure 6B**). Cells expressing IFI16-NLS mutant were more resistant to IAV infection and IAV induced cell death than cells expressing an equivalent amount of WT-IFI16 **(Figure 6C)**. Additionally, IFNβ was also more in cells expressing IFI16-NLS mutant than the cells expressing WT-IFI16 (**Figure 6D, 6E**). In contrast, cells expressing WT-IFI16 could bind more IAV-RNA than cells expressing an equal amount of IFI16-NLS mutant **(Figure 6F)**. Previous reports suggest that IFI16 binds to viral DNA through two tandem repeats of hematopoietic interferon-inducible nuclear (HINa, HINb) domains and one pyrin domain. We made multiple IFI16 mutants lacking either pyrin, HINa, HINb, or both HIN domains. IFI16 mutant having just the HINb domain was found to restrict viral infection similar to wild type IFI16, indicating a significant role of HINb domain (**Supplementary figure 4A, 4B**). Collectively, these data suggest that IFI16 is recruited into the nucleus of infected cells, where it senses IAV-RNA through HINb domain and initiates inflammatory cell death responses. We also found that only IFI16, among other members of ALR’s (AIM2-like receptors), plays a vital role in the regulation of Influenza virus infection (**Supplementary figure 4C**).

**Figure 6.**
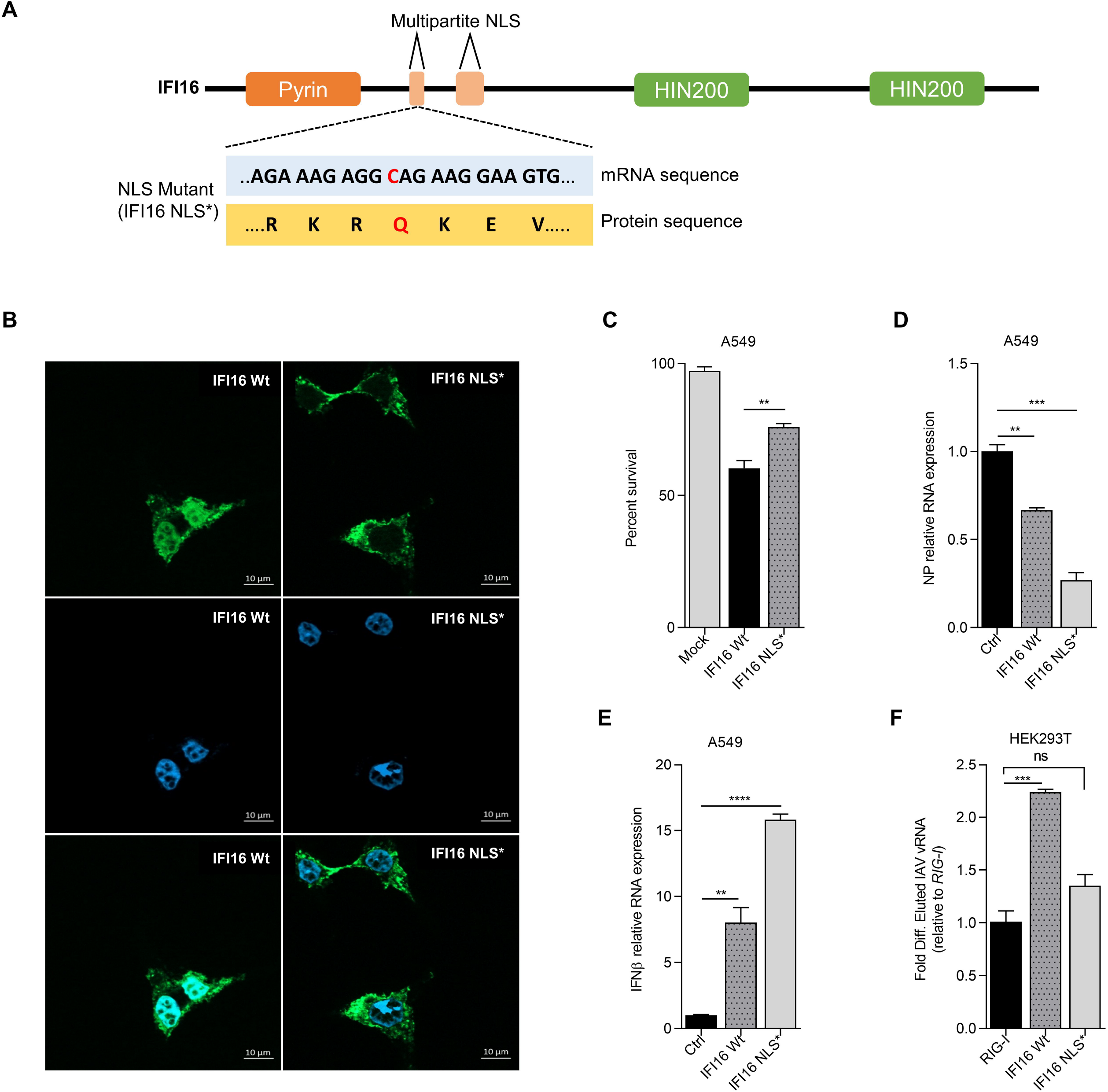
Nuclear localization of IFI16 is essential in restraining infection and sensing IAV genomic RNA. (A) Schematic showing the multipartite Nuclear Localization Signal (NLS) sequence of IFI16. The mutation made in the NLS sequence is marked in red. (B) GFP-tagged IFI16 and NLS mutant of IFI16 (IFI16 NLS*) were overexpressed in HEK293T cells. Cells were observed under a Microscope to find the IFI16 localization. (C) A549 cells were transfected with IFI16 Wt and IFI16 NLS* and infected with IAV (MOI 10) for 24 h. The cell viability was determined using the MTT assay. (D) (E) Relative expression of NP and IFNβ RNAs were measured in total RNA by qRT-PCR. (F) Flag-tagged IFI16 Wt, IFI16 NLS*, and RIG-I were overexpressed in HEK293T cells and then infected with IAV (MOI 10) for 24 h. RNA-IP was performed using Anti-flag M2 Affinity gel, and expression of viral RNA in the IP-ied sample was analysed by qRT-PCR using Input RNA for normalization.

## Discussion

PRRs sense conserved structures on invading pathogens and deploys various cellular and molecular components in antimicrobial response. Although our understanding of the innate immune system has exponentially increased in the past few decades. However, several questions such as how a limited number of germline-encoded PRRs orchestrate an array of immune responses against the spectrum of pathogens remains the most perplexing enigma in innate immunobiology. This report demonstrates that a single PRR can elicit an immune response against different classes of pathogen origin molecular patterns. IFI16 is a key viral restriction factor against DNA viruses. Upon sensing viral DNA, it induces pro-inflammatory cytokines through ASC-dependent inflammasome pathway and IFNβ through STING-TBK1-IRF3 signaling axis. For the first time, we have demonstrated that IAV genomic RNA serves as a ligand for IFI16. We also demonstrated that upon sensing, it induces IFI16-dependent programmed cell death to eliminate infected cells. In humans and other mammals, respiratory epithelial cells are the primary target of IAV infection, where being a lytic virus replicates in dying infected cells, causing desquamation in the respiratory tract of the host. Controlled, programmed cell death could be an effective host defense mechanism that suppresses cell-to-cell viral transmission and, subsequently, virus infection. Despite such an important host antiviral response, the exact molecular details driving programmed cell death upon viral infection remain unclear.

Caspase-1-dependent pyroptosis has been shown in respiratory and primary bronchial epithelial cells in *in vitro* condition, and similar exaggerated inflammatory response and cell death were also reported upon IAV infection *in vivo*. Our study has now identified a unique role of IFI16-mediated sensing of the viral RNA; it induces cell death in the form of apoptosis and pyroptosis. Another recent study has shown the RNA binding ability of IFI16 during Togaviridae family member, Chikungunya virus (CHIKV) infection, where IFI16 was found to restrict viral replication by binding to genomic RNA. In agreement with this, we found that IFI16 can bind to IAV genomic RNA as well. The two tandem repeats of hematopoietic interferon-inducible nuclear (HIN) domains of IFI16 are known to bind foreign DNA. Our results demonstrate that IAV RNA sensed by the IFI16 requires active replication of the virus in the nucleus. Consistent with previous findings of its DNA sensing ability, it indicates that the same domains that bind DNA can also bind RNA. Although the exact molecular mechanism, how a DNA sensing protein is binding to the RNA needs further investigation, this demonstrates the multi-role of PRRs in combating different types of virus infection.

Emerging RNA virus infections pose a significant threat to the socio-economic status of humankind. Finding drugs or vaccines for these emerging pathogens is highly challenging because of their high mutation rates. The ongoing Coronavirus pandemic associated vaccine and therapeutic discovery is a perfect example of this challenge. However, the mechanism of viral pathogenesis and innate immune strategies to block viral infection can pave the way for finding potential druggable targets. In this study, we provide evidence for the contribution of IFI16, a DNA sensor in sensing RNA from IAV and its downstream signaling results to cell death, suggesting that the multi-layer sensing system of IAV ensures all possible development of the appropriate and regulated antiviral state. Our study suggests IFI16, a member of ALR (AIM2-like receptors), as an innate immune sensor of IAV regulating antiviral responses. Insights gained from this study will improve our understanding of the mechanisms of regulation of IAV pathogenesis and lead to identifying more possible targets for therapeutic intervention.

## Acknowledgement

We thank Professor David Knipe for providing us IFI16 plasmid, R. Fouchier for providing the A/PR8/H1N1 reverse genetics system, Professor Yan Yuan for providing the pCMV3Tag1a plasmid. We thank IISER Bhopal for providing the Central Instrumentation Facility.

## Author Contributions

Conceptualization, S.M, and H.K; Investigation, S.M, A.S.R; Validation, S.M, A.S.R, A.K, A.R, P.K; Formal Analysis, S.M, A.S.R, A.K; Data Curation, S.M, and A.S.R; Writing-Original Draft, S.M, A.S.R and H.K; Writing-Review and Editing, A.S.R, A.K, and H.K; Project Administration, H.K; Funding Acquisition, H.K; Resources, H.K; Supervision, H.K.

