## Supplementary Figures for "Innate Immune sensing of Influenza A viral RNA through IFI16 promotes pyroptotic cell death"

### Supp Figure1

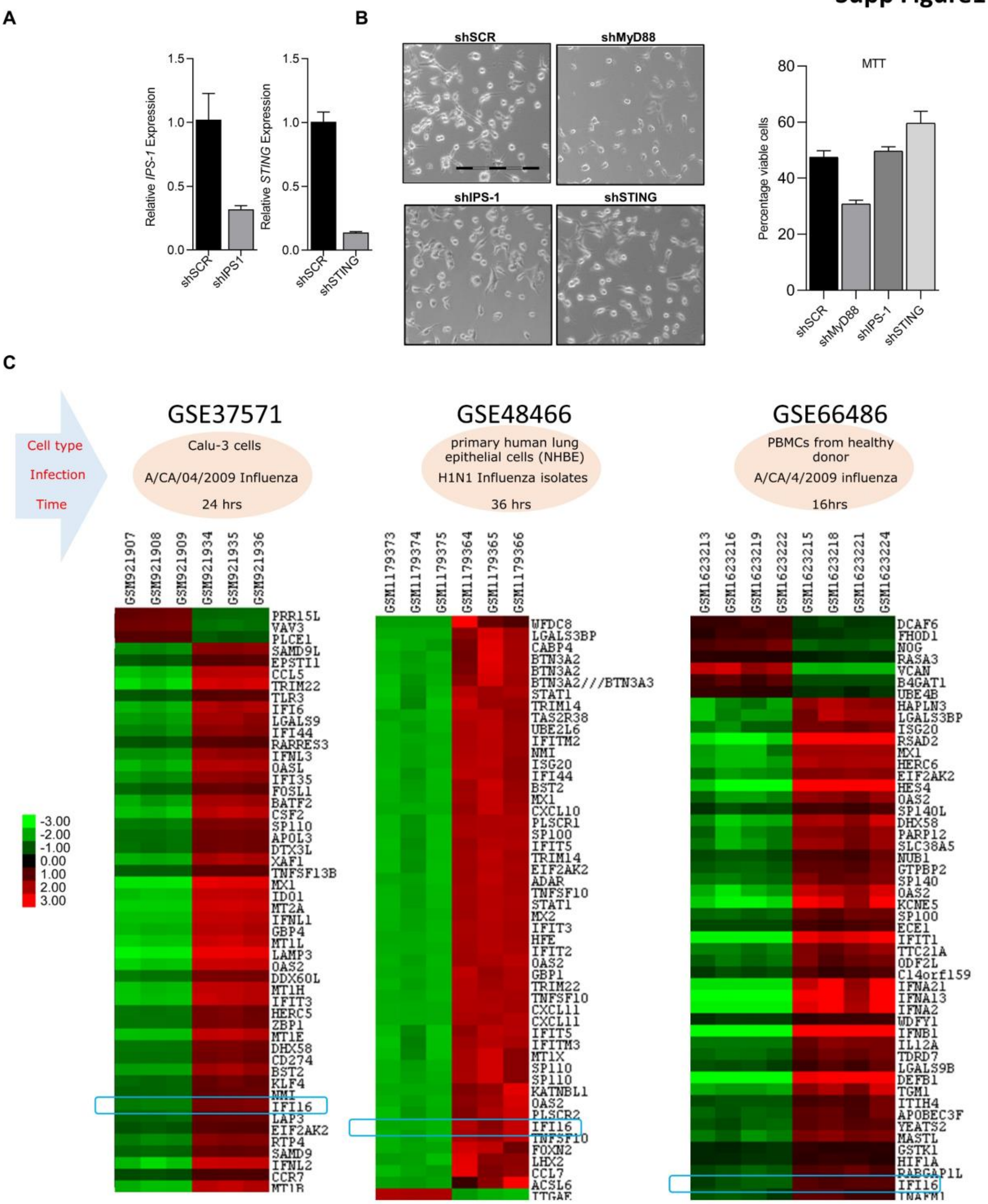


Supplementary Figure1- (A) A549 cells were transiently transfected with 2.0 μg of respective sh-clones of each indicated genes (shMyD88, shIPS-1, shSTING) or scrambled control (shSCR) for 48 h then infected with IAV (MOI 10) for 24 h. Knockdown efficiency was quantified using qRT PCR. (B) Microscopic images(10x) were taken, and the cell viability was determined using the MTT (Right panel). (C) Heat map analysis of the expression of nucleic-acid sensors and pro-inflammatory cytokines in IAV infection.

### Supp Figure 2

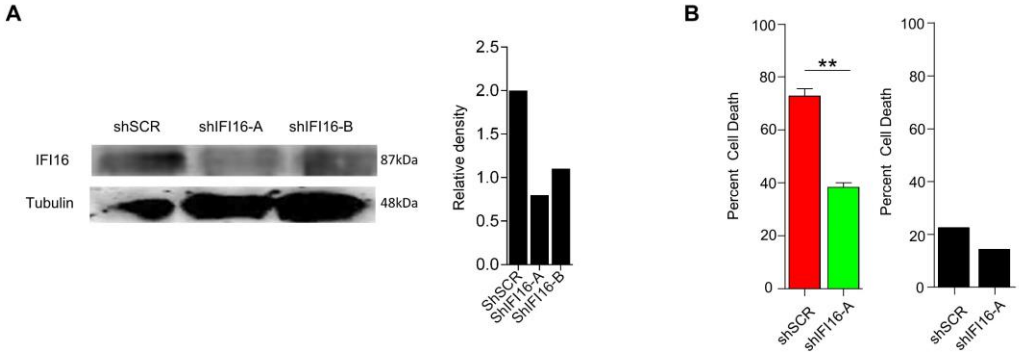


Supplementary Figure2- (A) A549 cells were transiently transfected with 2.0 µg of sh-clones of IFI16 (shIFI16) or scrambled control (shSCR) for 48 h then infected with IAV (MOI 10) for 24 h. Knockdown efficiency was quantified using Western Blot. (B)The cells were stained with Trypan blue, and the number of live and dead cells were counted using a hemocytometer.

### Supp Figure 3

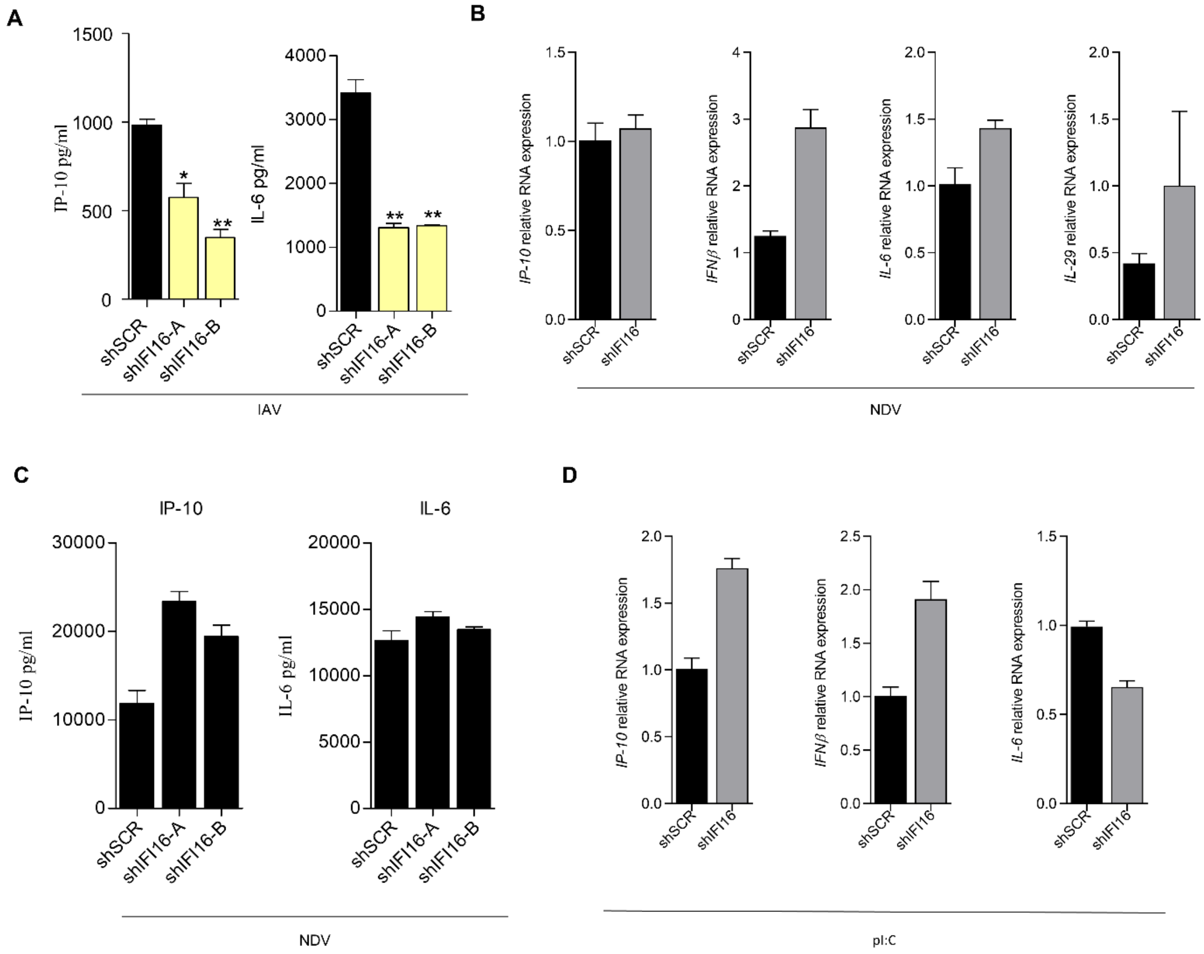


Supplementary Figure3- (A)A549 cells were transiently transfected with 2.0 µg of sh-clones of IFI16 (shIFI16) or scrambled control (shSCR) for 48 h then infected with IAV (MOI 10) for 24 h. Elisa for IL-6 and IP-10 was done from the supernatant. (B) A549 cells were transiently transfected with 2.0 µg of sh-clones of IFI16 (shIFI16) or scrambled control (shSCR) for 48 h then infected with NDV (MOI 2) for 24 h. Expression of type I interferon, type III interferon, and pro-inflammatory responses were analysed using qRT PCR. (C) Elisa for IL-6 and IP-10 was done from the supernatant. (D) A549 cells were transiently transfected with 2.0 µg of sh-clones of IFI16 (shIFI16) or scrambled control (shSCR) for 48 h then transfected with 1.0 µg of poly:IC. Expression of type I interferon and pro-inflammatory responses were analysed using qRT PCR.

**Supp Figure 4**

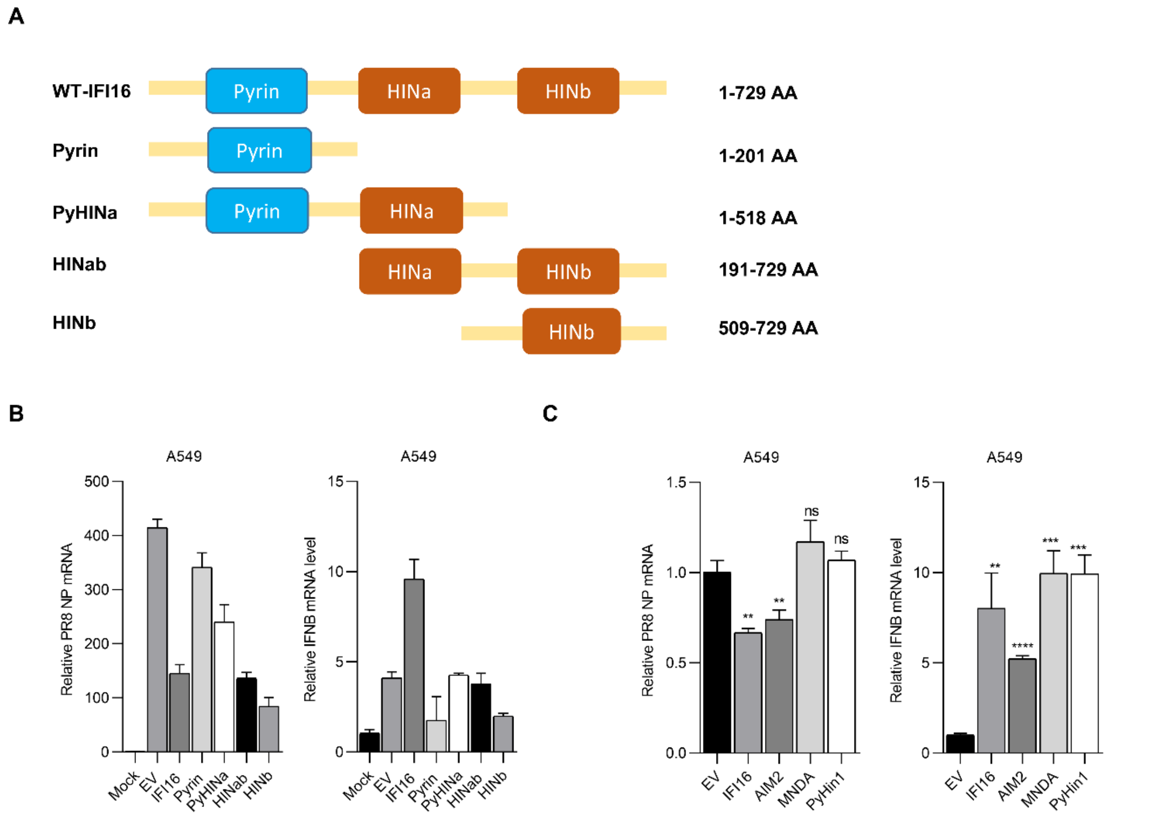


Supplementary Figure4- (A) Schematic showing the different domain mutants of IFI16. (B) A549 cells were transfected with plasmids expressing full length IFI16 or different domain mutants of IFI16 or the empty vector backbone (EV) and infected with IAV (MOI 10) for 24 h. Relative NP RNA and type I interferon expression were measured in total RNA by qRT-PCR. NP and IFNβ RNA expression was normalized to a mock-infected sample. (C) A549 cells were transfected with plasmids expressing IFI16 or different ALR family members or the empty vector backbone (EV) and infected with IAV (MOI 10) for 24 h. Relative NP RNA and type I interferon expression were measured in total RNA by qRT-PCR.
